# Accessibility and Reproducible Research Practices in Cardiovascular Literature

**DOI:** 10.1101/2022.07.06.498942

**Authors:** Gabriel Heckerman, Eileen Tzng, Arely Campos-Melendez, Chisomaga Ekwueme, Adrienne L. Mueller

## Abstract

**Background:** Several fields have described low reproducibility of scientific research and poor accessibility in research reporting practices. Although previous reports have investigated accessible reporting practices that lead to reproducible research in other fields, to date no study has explored the extent of accessible and reproducible research practices in cardiovascular science literature.

**Methods:** To study accessibility and reproducibility in cardiovascular research reporting, we screened 400 randomly selected articles published in 2019 in three top cardiovascular science publications: Circulation, the European Heart Journal, and the Journal of the American College of Cardiology (JACC). We screened each paper for accessible and reproducible research practices using a set of accessibility criteria including protocol, materials, data, and analysis script availability, as well as accessibility of the publication itself. We also quantified the consistency of open research practices within and across cardiovascular study types and journal formats.

**Results:** We identified that fewer than 2% of cardiovascular research publications provide sufficient resources (materials, methods, data, and analysis scripts) to fully reproduce their studies. After calculating an accessibility score as a measure of the extent to which an articles makes its resources available, we also showed that the level of accessibility varies across study types (p = 2e-16) and across journals (p = 5.9e-13). We further show that there are significant differences in which study types share which resources. We also describe the relationship between accessibility scores and corresponding author nationality.

**Conclusion:** Although the degree to which reproducible reporting practices are present in publications varies significantly across journals and study types; current cardiovascular science reports frequently do not provide sufficient materials, protocols, data, or analysis information to reproduce a study. In the future, having higher standards of accessibility mandated by either journals or funding bodies will help increase the reproducibility of cardiovascular research.

**Funding:** Authors Gabriel Heckerman, Arely Campos-Melendez, and Chisomaga Ekwueme were supported by an NIH R25 grant from the National Heart Lung and Blood Institute (R25HL147666). Eileen Tzng was supported by an AHA Institutional Training Award fellowship (18UFEL33960207).

## Introduction

Previous studies have reported on the lack of detailed methodology, data, and code provided that would be necessary for individual studies or study sets to be considered replicable and reproducible (Filazzola & Cahill, 2021). In addition, scientific publications are often behind paywalls that only users affiliated with a larger institution are readily able to access, meaning that research funded by public money is often inaccessible to the general public without paying a fee. Scientific practices have been evolving over time: data from a study can be complex and require a large amount of storage space, methods can be extremely sophisticated and require specialized equipment, and data analysis can require complex algorithms and code. Journals often do not specify requirements for materials sharing, data sharing, analysis code sharing, and methodological information - all of which have been shown to be important to be able to reproduce or replicate a study (Hamra et al, 2019; Munafò et al 2017). Note that often there may be legitimate reasons for restricting accessibility, such as privacy concerns, restrictive company policies, and costliness of the process.

Although some journals and funding agencies have implemented policies to support the public dissemination of research, we know from recent studies in several disciplines that accessible and reproducible research practices are still far from the norm (Borghi et al, 2018; Walters et al 2019; Kemper et al, 2020; Sherry et al, 2020; Smith et al 2021). However, to date, no study has examined the degree of accessible and reproducible practices in cardiovascular science publications. Within the domain of cardiovascular science research, this type of analysis is extremely important for holding the latest cutting edge discoveries in cardiovascular science to a high standard, because of the impact on human health. We therefore investigate reproducible reporting practices using an adapted previously published screening process (National Academies Press, 2019).

With this study, we hoped to identify the prevalence of accessible and reproducible research practices in cardiovascular research. To this aim, we defined what information is necessary to recreate research from a published work (reproducible). In conducting this study, we combined and evaluated the results of the screening process in order to create a statistical understanding of the overall state of current cardiovascular literature and pinpoint trends based on journal or article type. We screened randomly selected articles published in 2019 in Circulation, the European Heart Journal and the Journal of the American College of Cardiology (JACC). We tested, both, that the simple majority, more than half of screened publications, would lack one or more specific reproducibility or replicability criteria, and that the vast majority, over 90% of screened publications, would lack one or more specific reproducibility or replicability criteria. In addition, we predicted that some study types (e.g. clinical trials) would satisfy significantly more accessibility criteria than other study types. We also predicted that some categories of accessibility criteria would be satisfied significantly more frequently than other categories both study types and across separate journals (e.g. materials availability vs analysis script availability). We also specified that a lack of specific requirements by journals regarding what information to provide would lead to variability in which accessibility criteria are satisfied.

## Materials & Methods

### Sampling Plan

To determine the prevalence of accessible and reproducible research practices within cardiovascular literature, data was gathered from a random selection of all published studies in 2019 in the following three leading cardiology journals (Opthof, 2019): Circulation, European Heart Journal and the Journal of the American College of Cardiology (JACC). To limit the scope of our analysis, we only included articles published in the year of 2019. This specific year was selected because it was the most recent full year not influenced by changes in reporting practices due to the COVID-19 pandemic.

The following pubmed search string was used to obtain the full list of articles screened: ((“Circulation”[Journal] OR “J Am Coll Cardiol”[Journal] OR “Eur Heart J”[Journal]) AND 2019/01/01:2019/12/31[Date - Publication]) NOT (Published Erratum[Publication Type]). This search string retrieved 2786 articles. This list was randomized and the first 400 were screened for our study, and researchers screened the listed articles sequentially. No blinding was involved in this study and each article was screened at least twice by separate individuals. Some researchers screened a higher proportion or articles than other authors, but each author screened at least 100 articles once. Any ambiguities identified during screening were resolved either through additional review, or through discussion among participating researchers authors to achieve consensus.

### Variables

Following a modified version of a previously established coding protocol for reproducible and accessible research practices (Iqbal, Wallach et al, 2016), the following criteria were screened during the evaluation process: type of reported study, pre-registration status, pre-registration status, protocol availability, materials availability, data availability, analysis script availability, conflict of interest status, and open access of the article. An article’s “accessibility score” was calculated as the fraction of several accessibility criteria results with the specific criteria satisfied out of the total possible to be satisfied for that specific study type. See **Table 1** for a list of screening criteria that contributed to the calculation of the accessibility score. Note that for papers without the ability to share Materials (e.g. Meta-analyses), the materials criterion was omitted from the accessibility score calculation. Our article coding form is derived from Iqbal, Wallach et al 2016 and can be found as qualtrics.qsf and word files in the pre-registration (Campos-Melendez et al, 2021).

**Table 1:** Criteria determining accessibility score and criteria-satisfying responses.

| Criteria Screened | Criteria-satisfying responses |
| --- | --- |
| How clearly stated was the study type? | Stated (e.g. editorial comment, clinical trial) |
| Does the article state whether or not materials are available? | Yes the statement says that the materials (or some of the materials are available).* |
| Can you access, download and open the materials files? | Yes.* |
| Does the article state whether or not data are available? | Yes - the statement says that the data (or some of the data) are available. |
| Can you access, download, and open the data files? | Yes. |
| Are the data files clearly documented? | Yes. |
| Do the data files appear to contain all of the raw data necessary to reproduce the reported findings? | Yes. |
| Does the article state whether or not analysis scripts are available? | Yes - the statement says that the analysis scripts (or some of the analysis scripts) are available. |
| Can you access, download, and open the analysis files? | Yes. |
| Does the article state whether or not the study (or some aspect of the study) was pre-registered? | Yes - the statement says that there was a pre-registration. |
| Can you access and open the pre-registration? | Yes. |
| What aspects of the study appear to be pre-registered? (select all that apply) | Hypotheses, Methods, AND Analysis Plan all available. |
| Does the article link to an accessible protocol? | Yes. |
| What aspects of the study appear to be included in the protocol? (select all that apply) | Hypotheses, Methods, AND Analysis Plan all available. |
| Does the article include a statement indicating whether there were any conflicts of interest? | Any of the following can be selected:<br>Yes - the statement says that there are one or more conflicts of interest.<br>Yes - the statement says that there is no conflict of interest. |
| Does the article include a statement indicating whether there were funding sources? | Any of the following can be selected:<br>Yes - the statement says that there was funding from a private organization.<br>Yes - the statement says that there was funding from a public organization.<br>Yes - the statement says that there was funding from both public and private organizations.<br>Yes - the statement says that no funding was provided. |
| Is the article open access? | Any of the following can be selected:<br>Yes - found via Open Access Button.<br>Yes - found via other means. |

\* This criteria was not included for publications for which it was not possible to share materials.

The four criteria defining repeatability - methods (protocol or preregistration), materials, data, analysis script availability - are each considered an accessibility category. Each individual criteria in these five categories can be either satisfied or not. For a study to be considered fully “replicable” all three of the following categories must be available: methods, materials, and analysis scripts. For a study to be considered fully “reproducible”, all three of the following categories must be available: methods, data, and analysis scripts. If an article is either partially replicable or partially reproducible it is considered “partially repeatable”. Definitions of replicability and reproducibility were set based on those introduced by the National Academies of Sciences, Engineering and Medicine (NASEM) (2019). We defined the ability to replicate a study as the attempt to obtain the same results as a study by collecting new data. Reproducing a study describes the attempt to obtain the same results as a study by re-analyzing its data.

### Analysis

Only articles that were fully screened and for which all ambiguities had been resolved, were included in the final analysis. Papers written in a language other than English were also excluded, as well as any articles for which we could not access the full text. Articles without empirical data (e.g. review articles, commentaries, editorials), were evaluated for author location, language, conflict of interest, funding statements, and public access of the article itself, but were otherwise not screened for accessibility criteria.

To determine how frequently published cardiovascular research reports provide sufficient resources to replicate or reproduce their studies, we quantified the fraction of papers that are either partially replicable or partially reproducible - defined as partially repeatable. For all accessibility criteria, we calculated the proportion of papers in our dataset that exhibit each possible outcome. E.g. For the accessibility criterion “How does the statement indicate the materials are available,” we calculated the proportion of papers that state that materials are available through an online third party repository, a personal or institutional webpage, supplemental information hosted by a journal, or upon request from authors. We also calculated the distribution of accessibility scores across our full dataset and further collected data regarding the location of the corresponding author. We use standard deviations to show the variability of the data.

To test for inconsistent, or variable, levels of accessibility, we examined our data using multi-way ANOVAs. Our independent variables included “study type” and “journal” and our dependent variable was the accessibility score. For the hypotheses addressing differences between the types of studies, journals, and accessibility scores using multi-way ANOVA, p-values < 0.05 were considered to be statistically significant.

To evaluate the relationship between accessibility criteria (Materials, Methods, Data, Analysis Code) and study types we performed several sequential Chi square tests of proportions. We compared the proportion of papers that did and did not allow access to those four categories of information and resources, across all four categories. We also used chi square tests to further compare the proportion of papers that provided access to each category of information and which study types shared more or less of a specific type of information compared to other study types. We considered p-values <= 0.05 to indicate significant differences in proportions. We used Bonferonni correction to adjust the p-value for the number of tests performed. Note that this analysis differs slightly from that proposed in our pre-registration: Chi square Automatic Interaction Detection (CHAID).

### Pre-registration

This study was pre-registered through the Open Science Foundation and can be accessed at https://doi.org/10.17605/OSF.IO/QFSTH (Campos-Melendez et al. 2021).

After obtaining our data, not all exploratory analyses documented in the preregistration were performed. We did not test the relationship between the articles described as accessible and those actually accessible due to the limited number of articles that exhibited large enough accessibility scores, nor did we perform textual analysis on the conflict of interest, funding statements, or the way resources were being shared. We also did not perform additional analyses with the accessibility crtieria organized as a hierarchy. We also deviated from the pre-registration by using sequential chi square tests instead of the Chi square Automatic Interaction Detection (CHAID).

### Materials, Data, and Analysis Script Availability

All materials, data, and analysis scripts associated with this study are available on the Open Science Foundation website at https://doi.org/10.17605/OSF.IO/FUDKA (Tzng et al, 2021).

## Results

We screened a similar proportion of articles from each cardiovascular journal (Fig. 1C). Over half of the randomly selected articles from 2019 in the three top cardiovascular journals (247 out of 400 total) were review articles or other article types that contained no empirical data (Fig. 1A). Approximately half of the papers had easily-identifiable study types while the remaining papers required more substantial reading to determine their study type (Fig. 1B). Over half of the randomly selected articles (251) screened did not have a funding statement (Fig. 1D). Nearly all articles had a conflict of interest (COI) statement (357; Fig 1E), though notably over 10% did not. Additionally, over 95% of articles were publicly accessible (381 out of 400, Fig. 1F).

**Figure 1.**
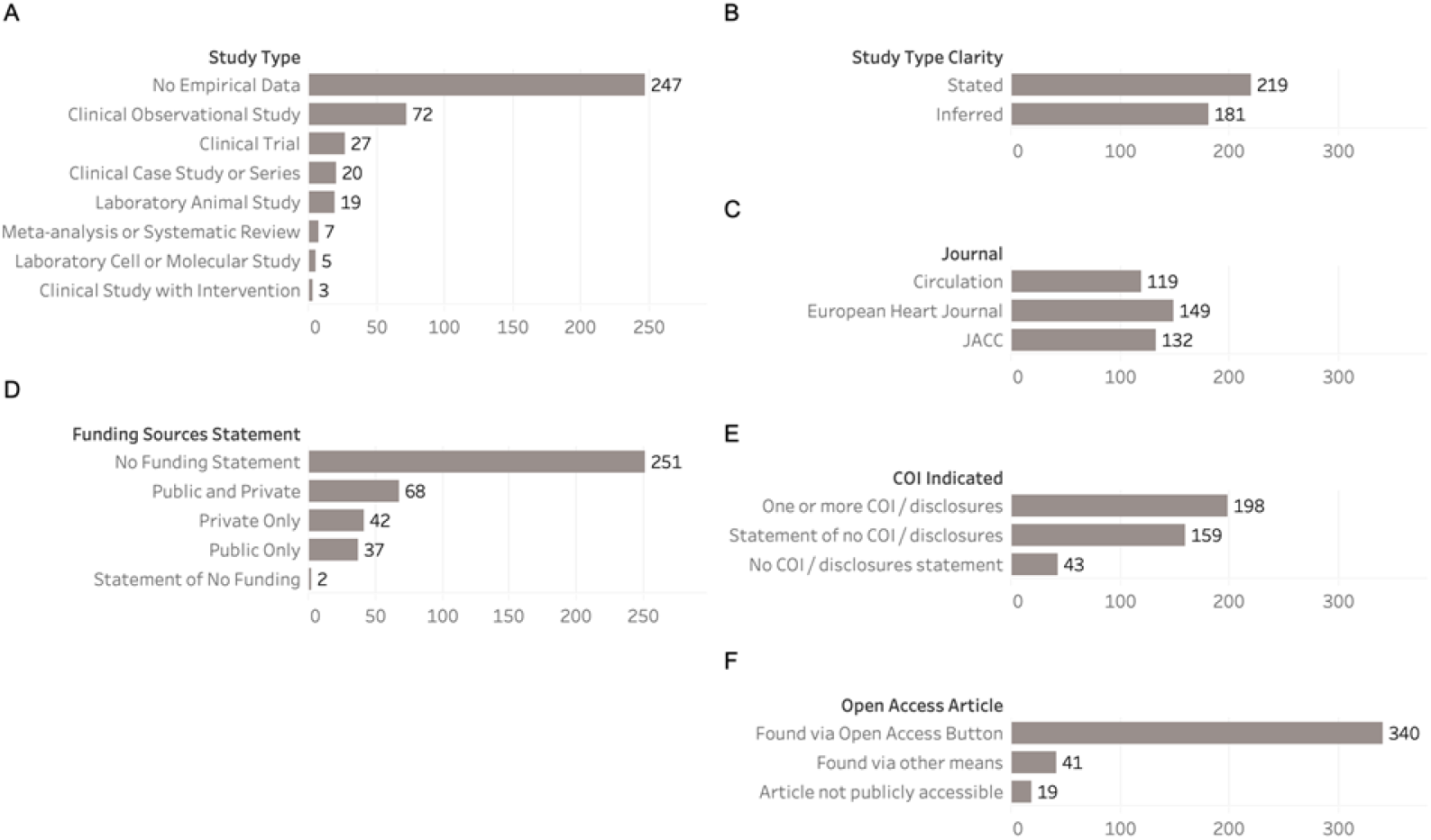
Summary of all screened publications on their (A) study type, (B) study type clarity, (C) journal in which it was published, (D) presence of funding source statement, (E) presence of conflict of interest statement, and (F) open access status.

### Cardiovascular research publications rarely make their resources available

Looking only at articles that included empirical research (153 out of 400 total articles), the majority had had no pre-registration statement (123) or no linked and accessible protocol (147; Fig. 2A). Of the papers that had a pre-registration statement, nearly all of them had an accessible and openable pre-registration (29 out of 30; Fig. 2A). Among articles that had an openable and accessible pre-registration or protocol, almost all articles included methods (29, 6), approximately half included hypotheses (12, 3), and less than half included analysis plans (7, 1; Fig. 2A). We define a publication to have all pre-registration aspects if hypotheses, methods, and analysis are pre-registered. Of the 153 screened publications that have empirical research, 14 had 1 out of 3 of the aspects, 11 had 2 out of 3 of the aspects, and 4 had all three aspects.

**Figure 2:**
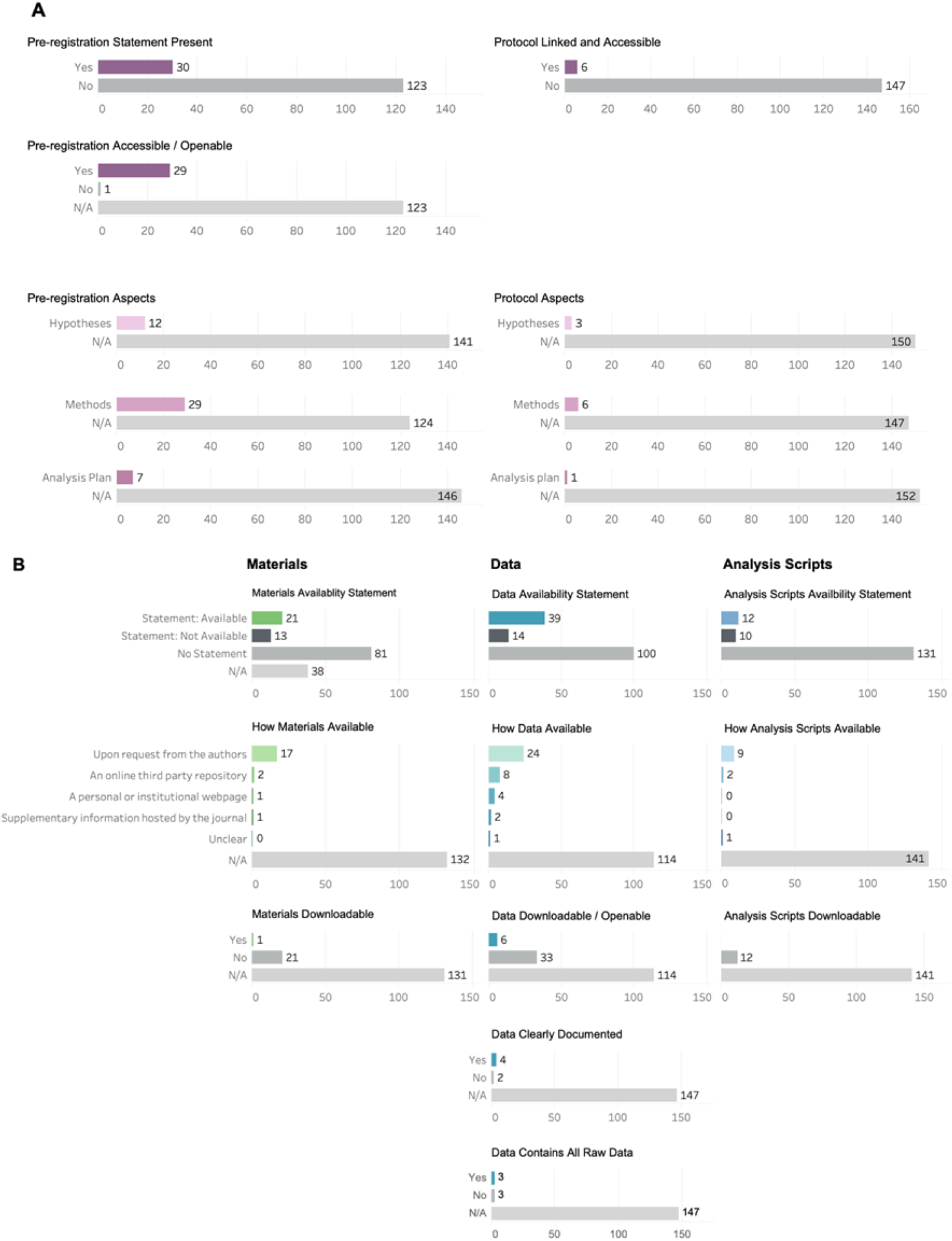
Summary of screened papers for pre-registration, protocol, material, data, and analysis script availability. Only study types considered empirical research in which materials are theoretically available were summarized. (A) Summary of the presence and accessibility of pre-registrations and protocols and the summary of components (hypotheses, methods, analysis plan) of pre-registration and protocol for papers that had a pre-registration or protocol statement. N/A represents papers that could not answer the criteria because data, materials, analysis plans were not available to begin with (top panel). (B) Summary of papers and material, data, and analysis script availability. For papers that had a statement, how materials, data, and analysis script were available, whether they were accessible, and whether it was clearly documented or present in its entirety are also summarized.

Only 14% of articles made their materials available (21/153), 26% of articles made their data available, and 8% made their analysis scripts available (Fig. 2B). Across the articles that stated materials, data, and analysis scripts were available, it was most common for them only to be made available upon request from the authors (Fig. 2B). Across the articles that stated materials, data, and analysis scripts were available, very few studies made the materials and data readily available for downloading or opening (1, 6) and no analysis scripts were readily downloadable (12 out of 12; Fig. 2B). However, the majority of data that was downloadable or openable was also clearly documented (4 out of 6; Fig. 2B).

The majority of cardiovascular publications are not readily replicable or reproducible. We predicted that the majority of publications will not provide sufficient resources to replicate or reproduce their studies. We tested both 1) that the simple majority, more than half of screened publications, will be lacking one or more specific replicability or reproducibility criteria, and 2) that the vast majority, over 90% of screened publications, will be lacking one or more specific reproducibility or replicability criteria. We found that 61.43% (94 out of 153) of empirical research papers were not partially reproducible and 68.62% (105 out of 153) were not partially replicable. Only 2 out of 153 empirical research papers were fully replicable, and only 2 were fully reproducible (Fig. 3). Therefore both the simple and vast majority of empirical research articles did not provide all the resources necessary to fully replicate or reproduce their studies.

**Figure 3:**
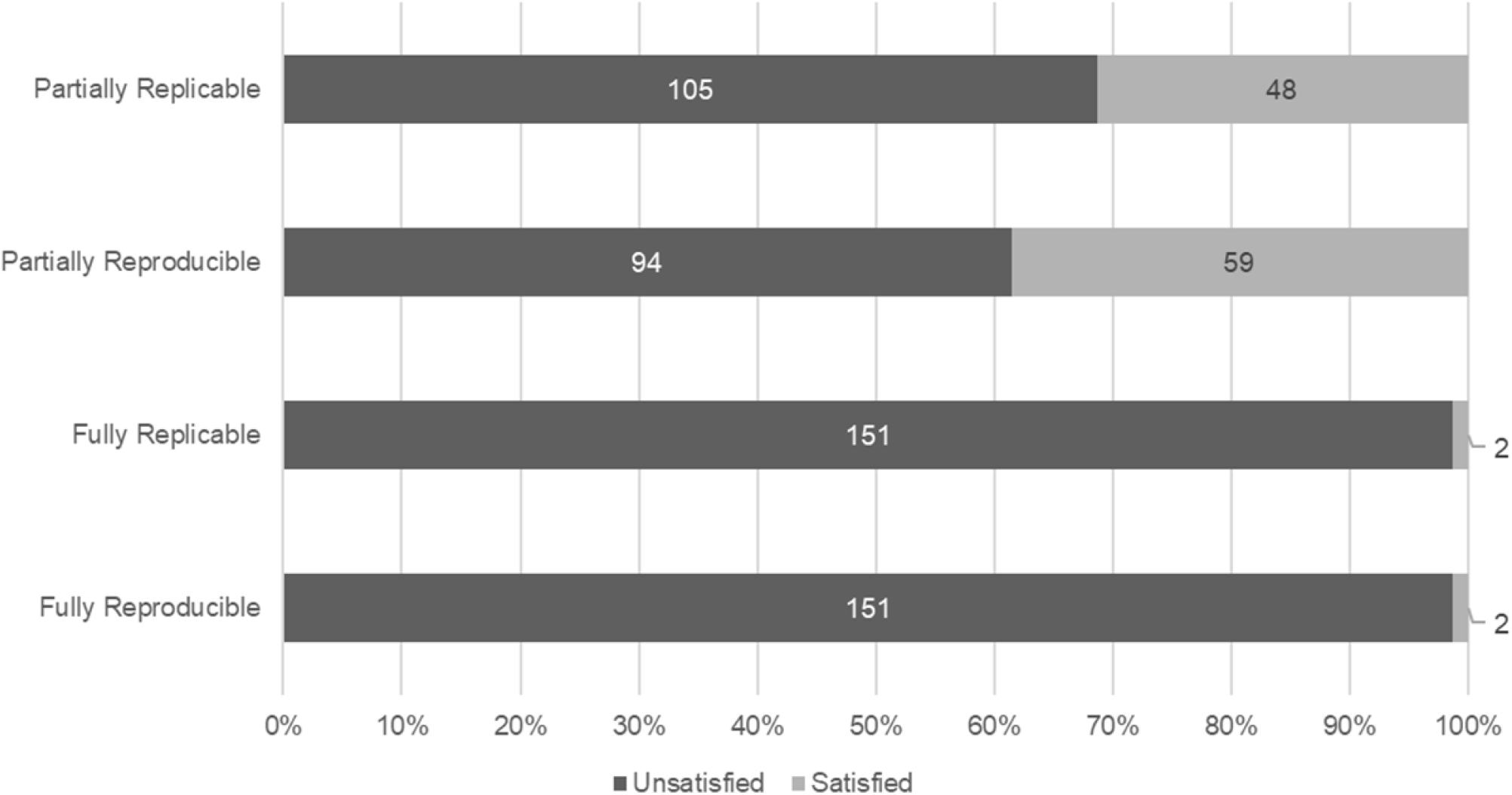
A summary of replicability and reproducibility of empirical research studies (n=153). An article is considered “partially replicable” if any of material availability, analysis script availability, and methods criteria are satisfied and “fully replicable” if all three criteria are satisfied. An article is considered “partially reproducible” if data availability, analysis script availability, and methods are satisfied and “fully reproducible” if all three criteria are satisfied.

### Accessibility Varies Across Journals and Study Types

With the exception of 3 articles, the entire data set exhibited low accessibility with scores lower than 0.5, with the majority of articles (99) having accessibility scores within the 0.1-0.299 range and 15 articles had scores below 0.1. No articles exceeded an accessibility score of 0.6 or greater (Figure 4A).

**Figure 4.**
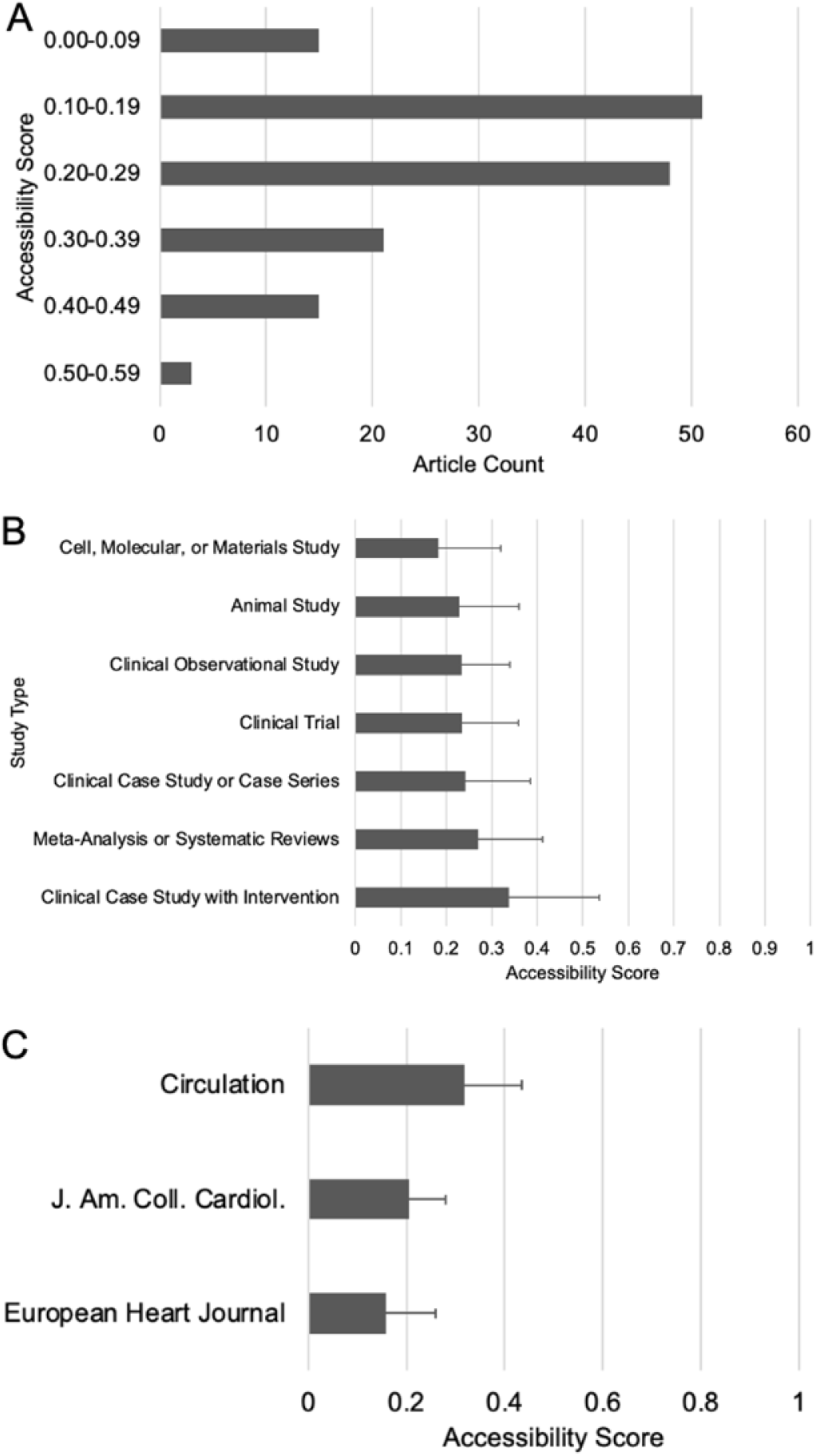
(A) Accessibility score distribution across the entire data set. The number of articles represented by each bar falls within the specified accessibility score range. All score fractions are based out of a total possible score of 1. No articles obtained an accessibility score fraction smaller than 0 or greater than 0.589. (B) Average weighted accessibility scores across all articles screened, based on study type. Error bars correspond to standard deviation of the collected data points. All scores are based on a total possible score of 1. (C) Average accessibility score by journal type. Screened articles were obtained from three different journals, including Journals of the American College of Cardiology (JACC), European Heart Journal from the European Society of Cardiology, and American Heart Association (AHA) Circulation. The standard error bars are indicative of the range of distribution of the obtained accessibility score fractions for each journal. All score fractions are based out of a total possible score of 1.

We also predicted that the level of accessibility will be inconsistent across several dimensions. Specifically, we predicted that average accessibility score would vary across study types and potentially also across journals, depending on the publisher’s requirements. For example, if clinical trials must be registered and open access, clinical trial publications will satisfy more of the article coding form criteria than other studies. We tested this hypothesis by calculating an accessibility score for every article: the sum of the satisfied screening criteria divided by the total possible satisfiable criteria. We then calculated the average and standard deviation in accessibility scores across study type (Fig. 4B). Overall accessibility scores were low across all study types, and standard deviations were large; indicating a high variability across individual publications. We found that clinical case studies had the average accessibility score (0.34), and cell, molecular, or materials studies had the lowest accessibility scores (0.18). Average accessibility scores were significantly different across study types (ANOVA, d.f. 28.4, F = 0.638, p = 2e-16).

We also predicted a significant difference in the number of accessibility criteria satisfied across journals due to differences in reporting policies. After computing the average accessibility score for all three journals, Circulation had the highest average accessibility score (0.32), followed by the Journals of the American College of Cardiology (0.21), then the European Heart Journal (0.16) (Fig. 4C). Following randomization, a different amount of articles were obtained from each journal type. Although the largest proportion of articles screened were obtained from the European Heart Journal with a total number of 149 articles only 29% of those were empirical research papers (43 out of 149). 38% of JACC articles (50 out of 132) and 51% of Circulation articles (60 out of 119) were empirical research for which we could calculate an accessibility score. Accessibility scores vary significantly across journals (ANOVA, d.f. 2, F = 34.17, p = 5.9e-13.) There was not a significant interaction between study type and journal (p = 0.194.)

### Categories of accessible resources varies across study types

We also predicted some categories of resources will be accessible significantly more frequently than other categories (e.g. materials availability vs analysis script availability). We found that there was a significant difference in the proportion of papers that satisfied specific accessibility criteria (*X*^*2*^ = 17.3011, p = 0.000613) We then further identified whether different study types exhibited differences in the proportions of accessibility criteria they satisfied. For example, whether there was a difference in the proportion of papers that have accessible data depending on study type. Note that we did not include Clinical Studies with Interventions, Meta-analysis or Systematic Reviews, or Laboratory Cell or Molecular Studies in this analysis due to their small sample size. For all four accessibility criteria (Materials, Methods, Data, and Analysis), we found a significant difference across study types (Table 2).

**Table 2:** P-values of chi square tests of study type for each category of resource.

|  | Yes | No | Study Type Chi Square p-value with Bonferroni Correction |
| --- | --- | --- | --- |
| Materials | 33 | 82 | 6.40E-04 |
| Methods | 32 | 121 | 7.20E-21 |
| Data | 53 | 100 | 2.24E-05 |
| Analysis | 22 | 131 | 2.25E-06 |

To determine specifically how study types differ in which accessibility categories they share, we performed multiple chi square tests with bonferroni correction, comparing each study type with every other study type for a given accessibility criteria (Fig. 5, Table 3). We found that a significantly higher proportion of laboratory animal studies stated that they would share materials than clinical observational studies. We also, as expected, found a significantly higher proportion of clinical trials shared their methods compared to any other study type. With regards to data sharing, we found that a significantly higher proportion of laboratory animal studies shared their data than either clinical case studies or clinical observational studies. Lastly, a significantly higher proportion of laboratory animal studies shared their data compared to clinical observational studies.

**Figure 5:**
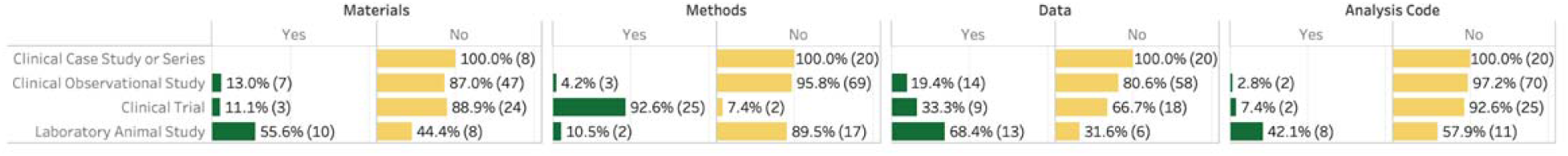
Proportion of papers with presence (yes) or absence (no) of specific accessibility criteria (Material, Methods, Data, Analysis Code) for specific study types. Presence is indicated in green, absence in yellow. For each value, both the percentage and the count for that category and study type are shown.

**Table 3:** P-values for chi square tests comparing the presence or absence of specific accessibility categories (Materials, Methods, Data, and Analysis Code) for each study type compared to every other study type (Clinical Case Study or Series, Clinical Observational Study, Clinical Trial, Laboratory Animal Study). P-values significant at an alpha level of 0.05 with Bonferroni correction are shown in red.

|  | Materials | Methods | Data | Analysis Code |
| --- | --- | --- | --- | --- |
| Clinical Case Study or Series vs Clinical Observational Study | 15.104 | 19.885 | 1.763 | 24.000 |
| Clinical Case Study or Series vs Clinical Trial | 18.946 | 4.91E-08 | 0.301 | 14.589 |
| Clinical Case Study or Series vs Laboratory Animal Study | 0.586 | 10.685 | 6.67E-04 | 0.102 |
| Clinical Observational Study vs Clinical Trial | 24.000 | 7.00E-16 | 5.616 | 15.340 |
| Clinical Observational Study vs Laboratory Animal Study | 0.018 | 14.538 | 0.003 | 1.94E-04 |
| Clinical Trial vs Laboratory Animal Study | 0.093 | 3.42E-06 | 0.978 | 0.346 |

### Accessibility scores vary across countries

We also collected data on corresponding author country and looked at the relationship between corresponding author country and accessibility scores (Fig. 6). Articles whose author’s indicated residence in Finland had the highest accessibility score with an average value of 0.41. One thing to note is the number of articles varied between each country, with the highest number of articles having corresponding authors from the United States of America (56). Most countries had a very limited sample size, meaning these scores may not accurately represent the level of average accessibility articles produced by corresponding authors from those nations.

**Figure 6.**
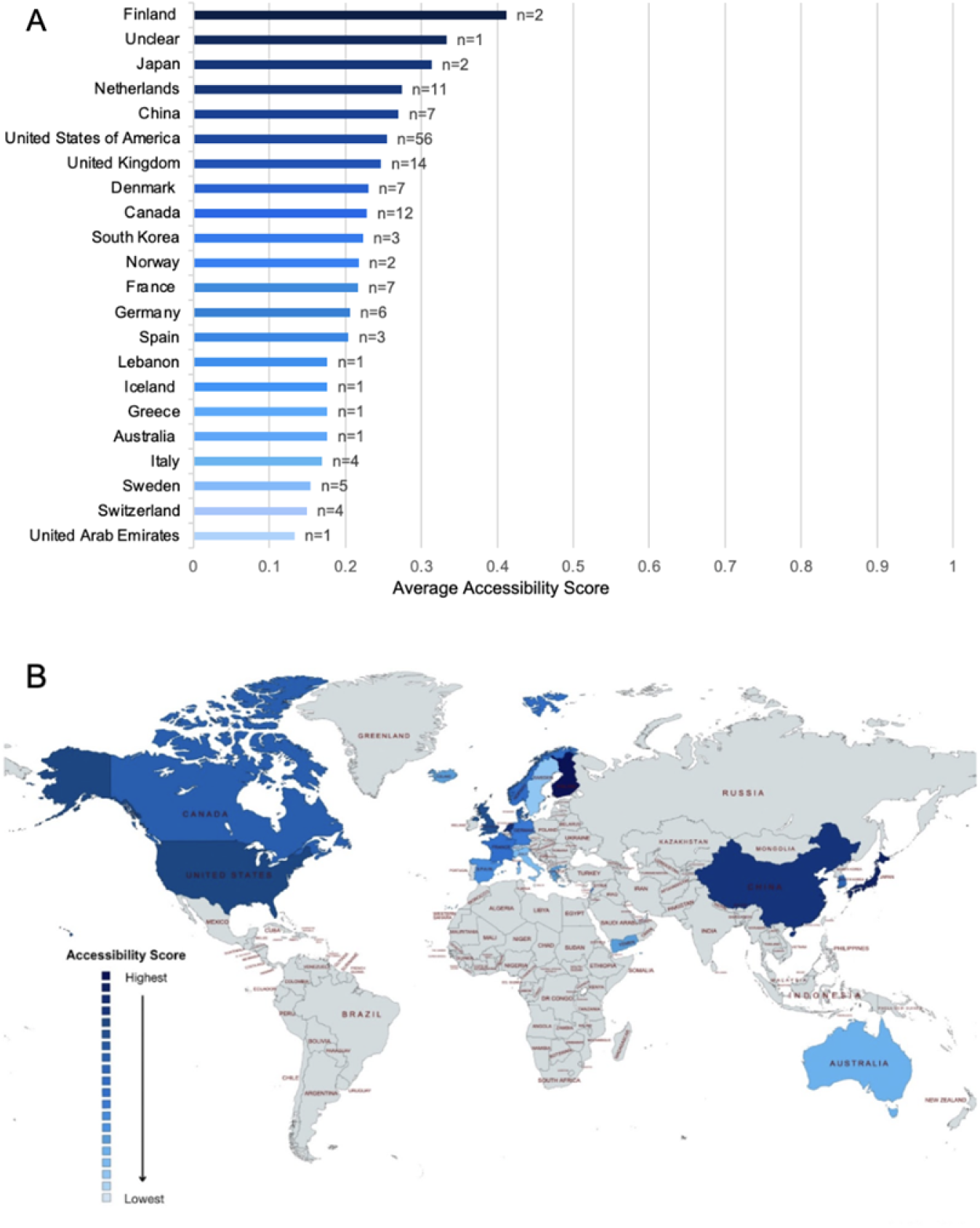
(A) Average accessibility score by corresponding author country of residence. Each bar represents the average accessibility score obtained by articles written by authors residing in the specified country. The number of articles used to calculate the average accessibility score for each country is indicated by the *n* value at the end of each bar. There were no average accessibility scores greater than 0.412 nor lower than 0. All accessibility score fractions are based on a total possible score of 1. (B) Worldwide distribution of accessibility scores based on author country of residence. Higher accessibility scores are depicted with darker shades of blue where lighter shades correspond to lower scores. Finland is colored with the darkest shade, indicative of its highest score; the lightest shade corresponds to the United Arab Emirates. The remainder of the countries had either no articles included in the screening process or only had articles without the empirical data necessary to calculate accessibility scores.

## Discussion

We reviewed 400 papers that were published in 2019 in three of the top cardiovascular research journals to determine how accessible, and therefore reproducible, their research was. Our intention with this study was not to disparage previous reports, but to share current practices with the cardiovascular field and identify ways to improve those practices to enhance reproducibility and transparency in the future. We hope to identify opportunities for journals and scientists to adapt their practices to further reproducible science, and encourage the use of higher standards, and consistent formats, among cardiovascular scientific literature.

In general, we found that the simple majority but not the vast majority of publications are lacking one or more of the resources (materials, methods, data, or analysis scripts) to replicate or reproduce a study. Only 2 out of 153, or less than 2% of papers, provided sufficient resources to fully replicate or reproduce their work. Although there were statistically significant differences in accessibility scores across study types and journals, overall, accessibility scores were consistently low.

### Publication Accessibility

In collecting data on the fraction of papers that were publicly accessible, we found that only 19 out of 400 articles were not publicly accessible (Fig. 1F). Although this is a fairly low fraction of inaccessible articles, a study’s publication is in many ways the most tangible output of the research: and having publicly accessible publication should be a minimum standard for research. The NIH’s public access policy has been instrumental in ensuring public access to published research reports, but reports funded purely through private sources are not under the same reporting mandate.

### Study Type

It is notable that over half of the 400 papers we randomly selected for screening were non-empirical research, e.g. reviews, editorials and commentaries. Although these emissions may be effective ways to communicate quickly and effectively with larger audiences, they may not go through the same rigor of review process. We also occasionally screened articles that were simplified summaries of the original research study, geared towards a lay audience. This type of article is valuable in that it makes research more accessible to a broader audience; however, because methods were absent and results were condensed, it was ambiguous whether the studies being reported in these articles had undergone full scientific review. Readers run the risk of assuming that these articles describe a full scientific story as opposed to a news highlight.

In addition, publications of replication studies, or studies that included a replication study, were virtually absent from our dataset (2 out of 153 empirical research publications, or 400 publications in total) - suggesting the field of cardiovascular research puts very little emphasis on replication work. Of concern, several studies also stated outright that their data was not available for replication: “The investigators will not make the data, methods used in the analysis, and materials used to conduct the research available to any researcher for purposes of reproducing the results or replicating the procedure.” Although there can be legitimate reasons for why data sharing is not possible, this statement provides no further explanation. This could lead to speculation of author motivations.

### Materials, Methods, Data, and Analysis Script Sharing

Although data was shared more frequently than methods, materials, or analysis scripts, there were still only a total of 6 out of 153 empirical research papers for which data was readily accessible to a reader. We acknowledge that authors will often have legitimate reasons for not being able to share data, resources, and more, including patient privacy. However, it has also been shown that patients are in general very willing to make their data available to further research (Seltzer et al, 2019; Kim et al, 2019) We advocate for research studies actively seeking consent from human subject participants to make their data available and if that consent cannot be obtained to specify that specific reason for not sharing data.

Although basic research studies more frequently shared materials, data, and analysis scripts than clinical trials, that sharing was frequently “upon request from the authors,” which previous studies have shown to be hit-or-miss in terms of yield (Hardwicke et al, 2018). Clinical trials had by far the most consistent availability of a resource category: methods, in the form of pre-registrations. Because pre-registrations are mandated by the FDA in order to conduct a clinical trial; our results suggest that resource-sharing requirements by funding or approval agencies are effective means of changing practices.

### Conflict of Interest (COI) and Funding Statements

Publications are a critical form of communicating research, and the content and results of publications are often prioritized to the point that COI and funding statements are easy to overlook. As a community, we are often more interested in the results described in publications than the process behind getting them. Personal interests and finances are undeniably a part of experimental integrity and can impact experimental design and therefore COI and funding statements should be given as much attention as other components of the publication.

Although it was beyond the scope of this study to perform a rigorous analysis of conflict of interest (COI) and funding statements, our screening process did reveal numerous cases of ambiguous COI and funding declarations. In Table 4, we capture both problematic and good examples of COI and funding statements. For example, many COI and funding statements were vague in that they lacked details on how different funders of interests specifically influenced the study, which is important for interpretation of the results. As another example, we also identified “Disclosures: None” as a problematic statement, because it could be interpreted as the authors having no disclosures, or, that the authors declined to list their disclosures. The relationship between funders and COIs is also ambiguous. Authors frequently listed funders and declared “no conflict of interest;” however, funders frequently do have an interest in and impact on the study and therefore represent a conflict of interest. If nothing else, it is in the interest of researchers to produce compelling results to maintain good relations and receive future funding from agencies, even if they were funded solely through public organizations.

**Table 4:**
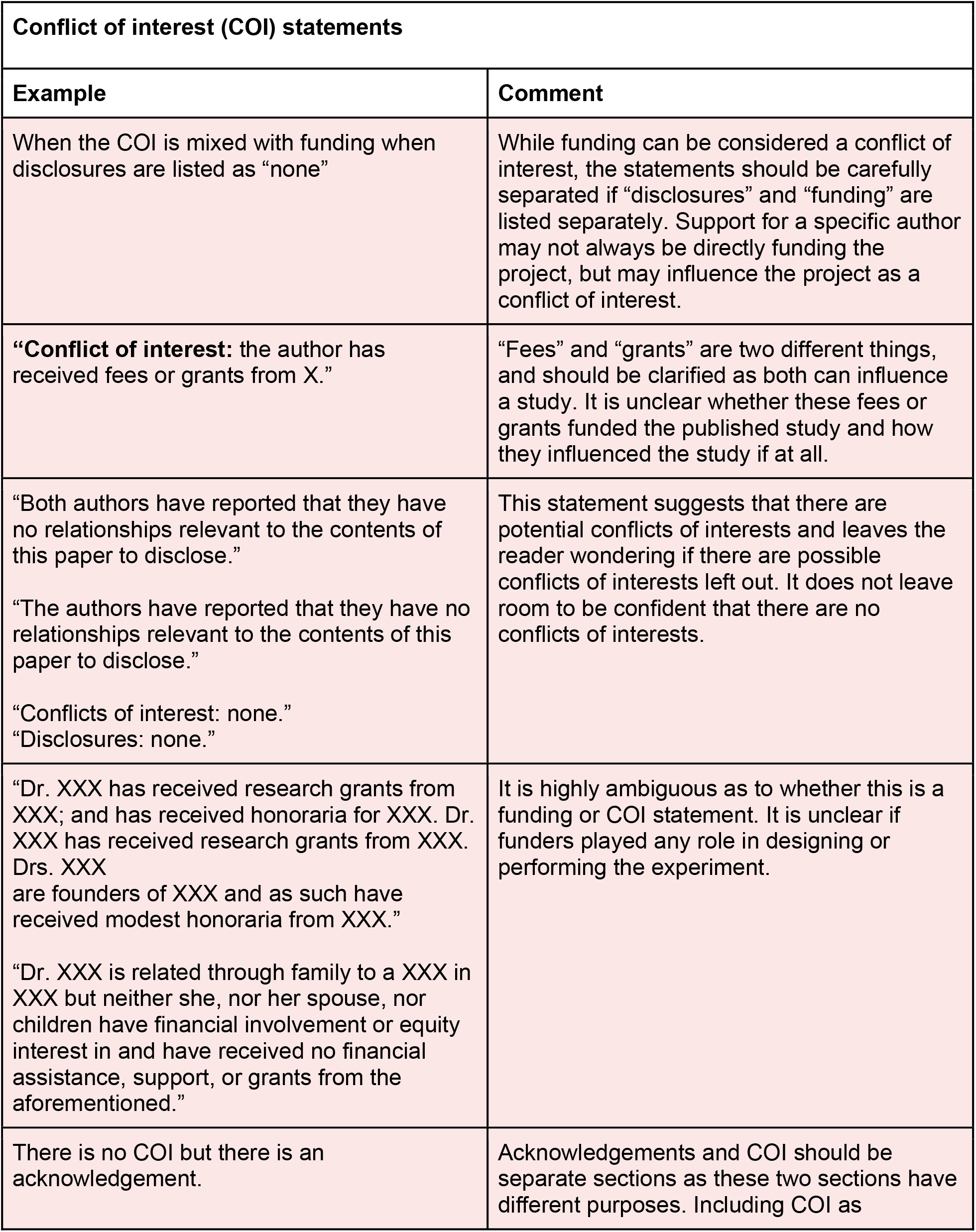
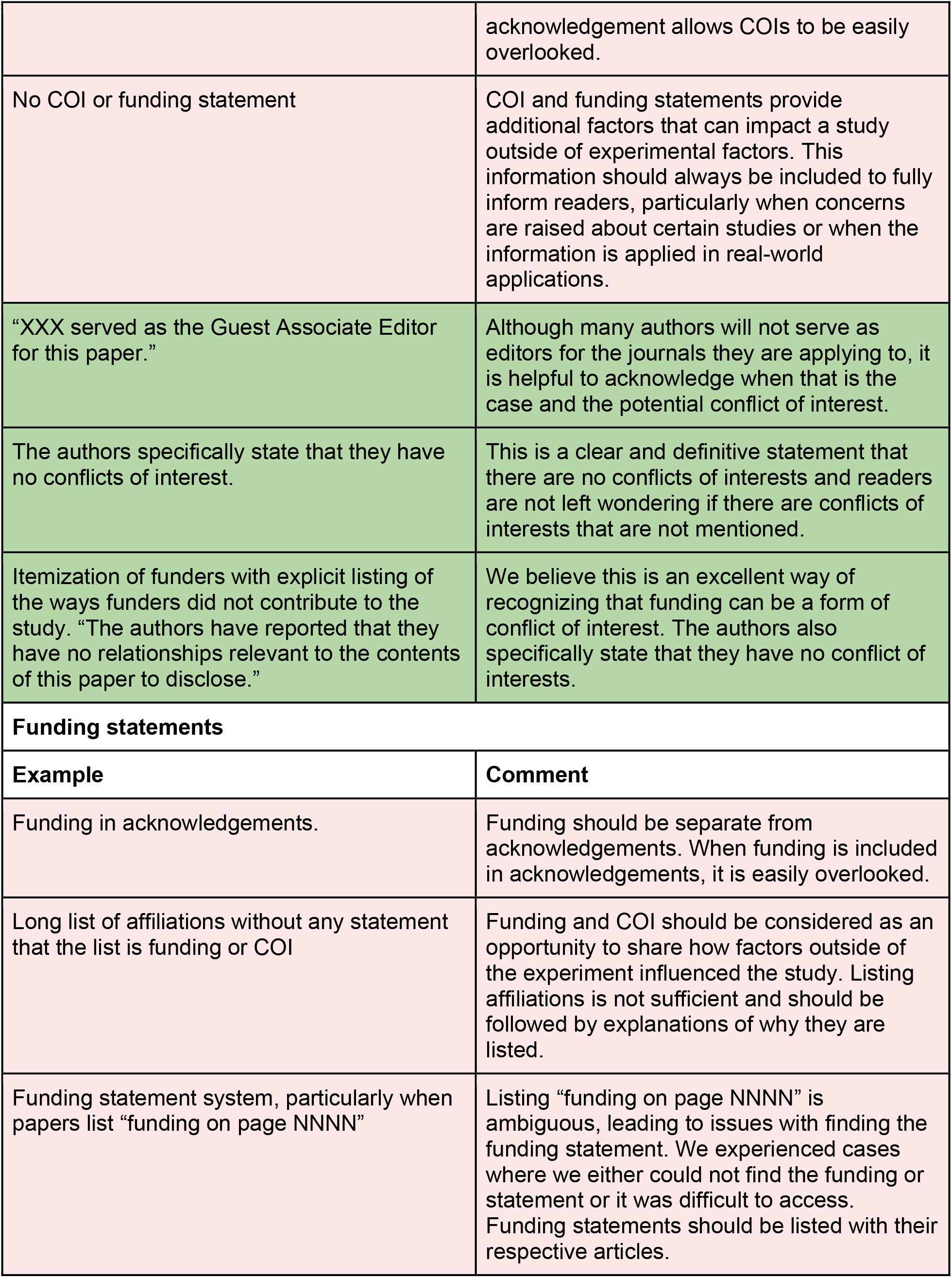
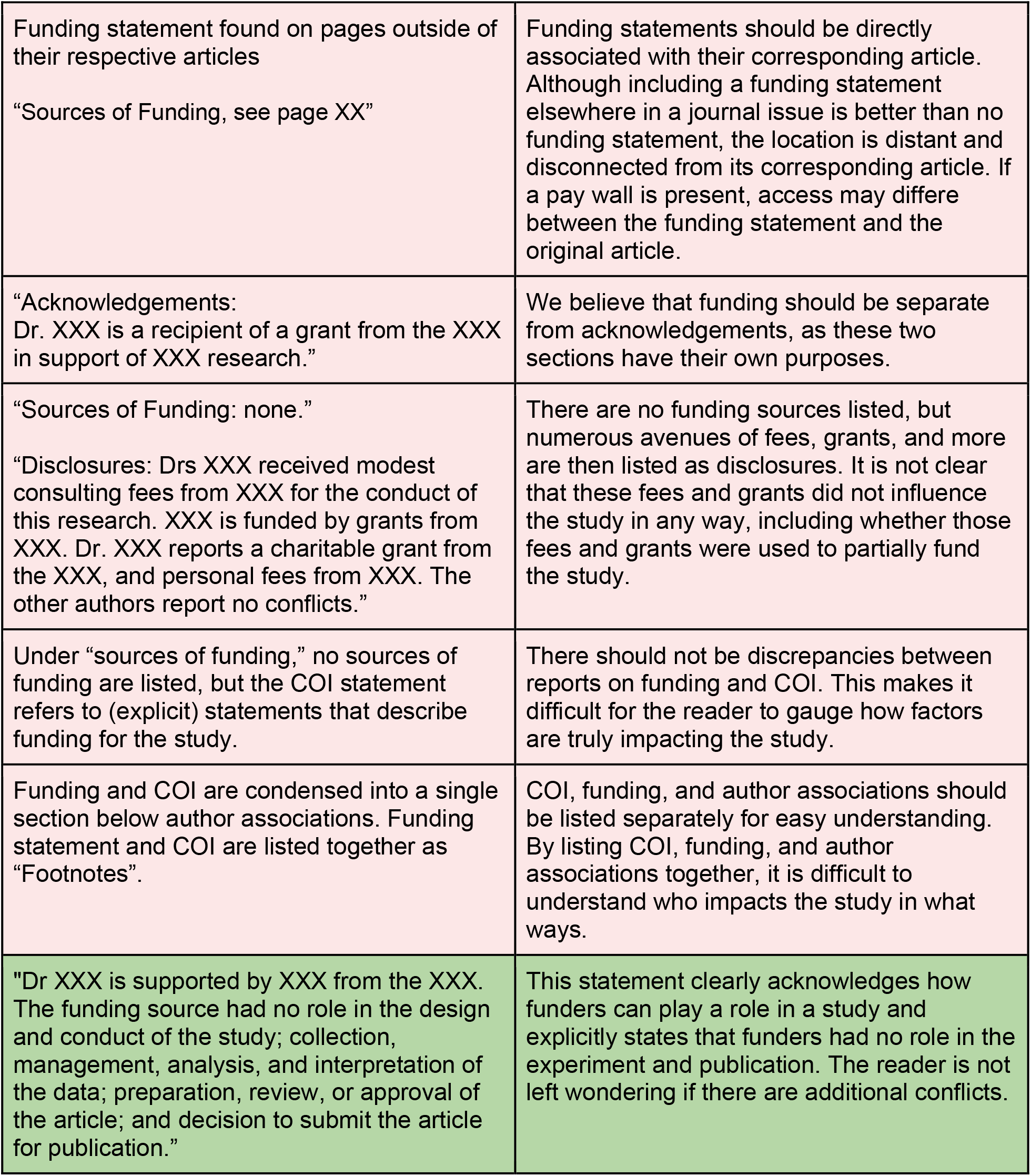
A summary table of problematic and informative COI and Funding statement examples that we repeatedly came across. Ambiguous or problematic statements are in yellow and clear statements are in green.

To avoid these issues, we advocate that journals require more complete and standardized COI and funding information that recognizes their overlap, including: a clear description on how each funding entity supported the work including whether they influenced the experiments, analysis, or dissemination of the work. We recommend that journals require authors to complete a consistent and transparent funding and COI questionnaire or decision tree that is clearly and prominently associated with the article.

### Authorship

Although we did not have sufficient articles from different countries of corresponding authors to determine whether there was a significant difference in accessibility practices across countries, we did see a wide range in accessibility score values. In terms of reporting, it was in general very straightforward to identify the corresponding author of an article. However, sometimes corresponding authorship was ambiguous in that no contact information was given, or was listed as, “Published on behalf of […]. All rights reserved. © The Author(s) 2019. For permissions, please email: […].” In these circumstances, it is unclear iwho the corresponding author is and who is ultimately responsible for the research.

### Limitations

Although we attempted to achieve completeness and consistency in screening by ensuring each article was screened by two separate individuals, we acknowledge we may have missed or misinterpreted statements in screened articles. For example, in capturing the type of funding for a study as public, private, or a combination of both, we may have misidentified the funding type for some organizations, especially for foreign funding bodies. Note that regular readers of publications will face the same challenges.

Additionally, we did not anticipate the high proportion of studies in our dataset of 400 articles that did not include any empirical research. Therefore, the number of studies upon which we could perform accessibility analyses on was reduced to less than half of our total pool of papers. Although we had sufficiently high numbers of papers to perform the majority of our analysis, having a larger sample size would have allowed more extensive comparisons across study types and accessibility categories.

This study also only used data from some of the highest ranking cardiovascular research journals. These journals are likely to have higher reporting standards than other journals in the field, so our results are likely to overestimate the reporting and sharing practices of publications in cardiovascular research in general.

### Future Work and Conclusion

The data collected from 400 screened papers provides numerous future directions for not only exploratory analysis on the existing data set but also for new projects assessing accessibility and reproducibility of scientific literature. The accessibility scores we calculated are a rough, quantitative estimate of an article’s actual accessibility and further work is required to more fully describe how accessible and reproducible cardiovascular literature is. For example, future work could identify which criteria are the biggest needs for the field, and then evaluate the quality of an article’s accessibility by weighting those criteria or organizing the criteria into a hierarchy of importance. Future studies could also investigate text excerpts describing how resources are or are not being made available to determine the causes that promote or undermine accessible research practices.

Our study shows that there is a high degree of variability in the resources cardiovascular research publications make available - across study type, journal, and country. Universally, however, publications almost never provide sufficient materials, protocol information, data, or analysis scripts for another group to fully replicate or reproduce their work. When federal policies mandate the open research practices, such as ensuring publications are publically accessible or clinical trials are pre-registered, they are adopted. To ensure the highest caliber of research in the future, we urge journals and funding agencies to require higher standards in their materials, protocol, data, and analysis script sharing - regardless of the study type or the funding source.

